# Cell autonomous polarization by the planar cell polarity signaling pathway

**DOI:** 10.1101/2023.09.26.559449

**Authors:** Alexis T Weiner, Silas Boye Nissen, Kaye Suyama, Bomsoo Cho, Gandhy Pierre-Louis, Jeffrey D Axelrod

## Abstract

As epithelial cells polarize in the tissue plane, the Planar Cell Polarity (PCP) signaling module segregates two distinct molecular subcomplexes to opposite sides of cells. Homodimers of the atypical cadherin Flamingo form bridges linking opposite complexes in neighboring cells, coordinating their direction of polarization. Feedback is required for cell polarization, but whether feedback requires intercellular and/or intracellular pathways is unknown. Using novel tools, we show that cells lacking Flamingo, or bearing a homodimerization-deficient Flamingo, polarize autonomously, indicating that functional PCP subcomplexes form and segregate cell-autonomously. Furthermore, we identify feedback pathways and propose an asymmetry amplifying mechanism that operate cell-autonomously. The intrinsic logic of PCP signaling is therefore more similar to that in single cell systems than was previously recognized.

## Introduction

The planar cell polarity (PCP) signaling pathway polarizes individual cells and coordinates polarity between cells along an axis parallel to the tissue plane^1^. The mechanism by which PCP polarizes cells has been intensively studied in *Drosophila*, and is substantially conserved from flies to humans^2^. Medically important developmental defects and physiological processes in vertebrates are also under control of PCP signaling, motivating interest in understanding the underlying mechanisms^1,3^. Major advances in understanding molecular mechanisms have come from leveraging of genetic approaches only possible in *Drosophila*, and evident conservation reinforces the utility of flies as a model system.

Proteins in the core PCP module, including the seven-pass atypical cadherin Flamingo (Fmi; a.k.a. Starry night)^4,5^, the serpentine protein Frizzled (Fz)^6,7^, the 4-pass protein Van Gogh (Vang; a.k.a. Strabismus)^8,9^, and the cytosolic/peripheral membrane proteins Dishevelled (Dsh)^10,11^, Diego (Dgo)^12^, and the PET/Lim domain protein Prickle (Pk)^13^ (reviewed in ref.^1^) segregate within cells to form distinct complexes on opposite sides of the cell (Fig. 1A,B). A subcomplex containing Dgo-Dsh-Fz-Fmi and an oppositely localized Fmi-Vang-Pk subcomplex communicate their presence and identity to the adjacent cell via intercellular bridges formed by Fmi=Fmi homodimers, thereby coordinating polarity between cells^14–18^. Any of a number of proposed subtle biasing signals can determine the direction of polarization, and bistability in the core system along with coordination between cells is proposed to reinforce the response^19^. Both the localized Dgo-Dsh-Fz-Fmi subcomplex and the oppositely localized Fmi-Vang-Pk subcomplex appear to direct cytoskeletal regulators to control morphological outcomes. In the wing, they determine the position and orientation of emerging trichomes (a.k.a. prehairs)^20,21^.

**Figure 1.**
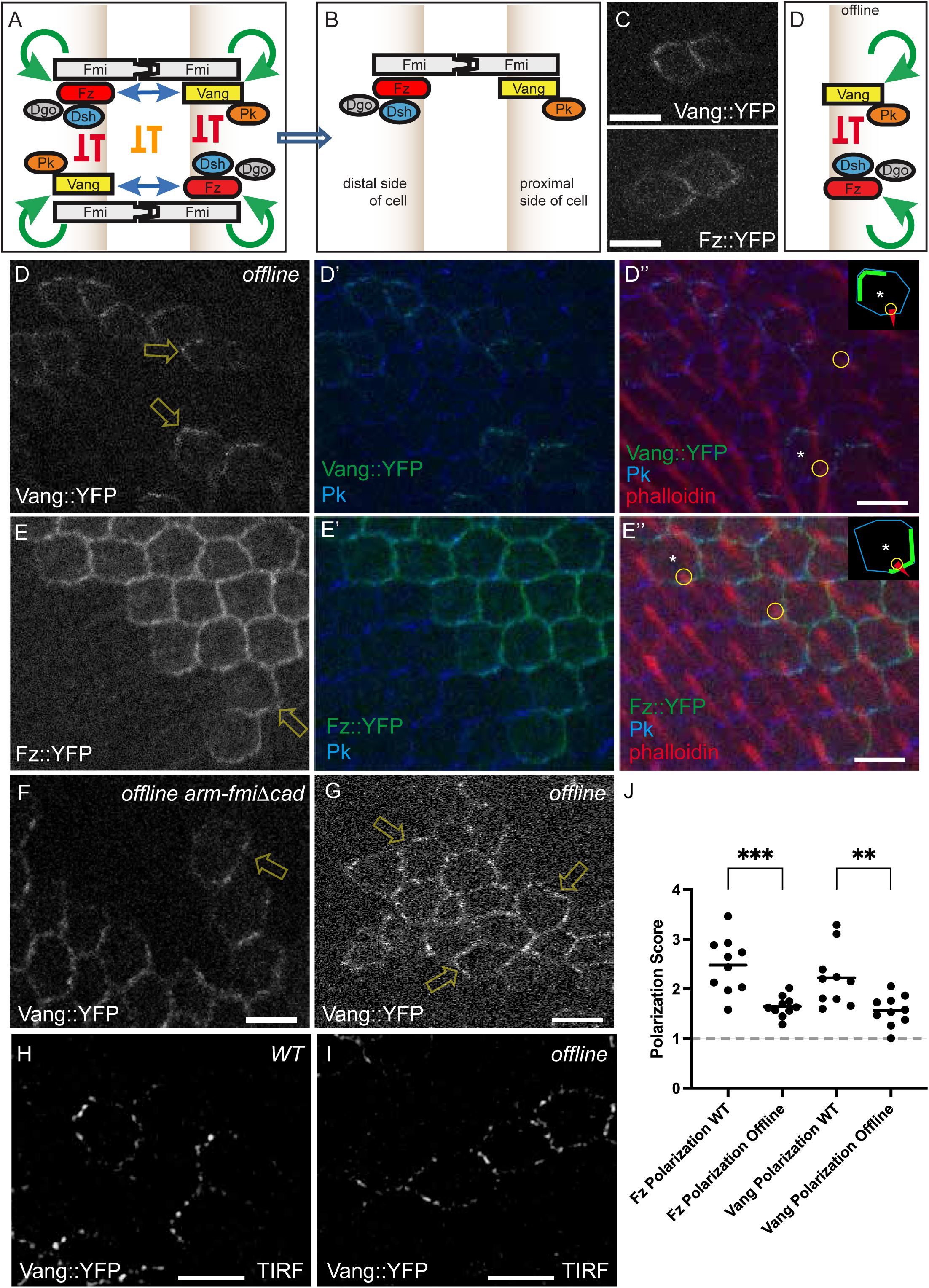
Offline cells polarize. A,B. Schematic of PCP complex composition and organization. Asymmetric Fmi bridges linking Fz-Dsh-containing sub-complexes to Vang-Pk-containing sub-complexes (A) sort themselves to yield predominantly one sub-complex on one side of the cell and the other sub-complex on the opposite side of the cell (B). Green and blue arrows represent hypothesized positive feedback pathways and red and orange arrows represent hypothesized negative feedback pathways. Blue and orange pathways require intercellular communication whereas green and red might work cell-autonomously. In all schematics, the depicted proteins are near the apical end of the lateral cell surfaces and intact complexes form intercellular junctions C. Vang::YFP and Fz::YFP clones in a wild-type background. For merged images showing clones with prehairs, see Supplementary Figure 2. In this and all panels throughout the manuscript, images are displayed with distal to the right and anterior up. D-D’’. Vang::YFP clones in 34 hr APF offline (*fmi^null^*) wings reveal asymmetric segregation as occurs in wildtype wings. N=10 individual wings from different animals. The direction of polarization deviates from the proximal-distal axis (arrows) and corresponds to the direction of prehairs (D,D’). Prehairs arise on the side of the cell opposite to the Vang::YFP crescents as occurs in wildtype (D’’; circles). The inset here and in panel E trace the cell marked with an asterisk, showing the cell outline, the crescent of YFP signal, and the location and orientation of the prehair. E-E’’. Fz::YFP clones in a 34 hr APF offline wing. N=10 individual wings from different animals The direction of polarization deviates from the proximal-distal axis (arrows) and corresponds to the direction of prehairs (E,E’). Prehairs arise on the same side of the cell as the Fz::YFP crescents as occurs in wildtype (E’’; circles). F. Vang::YFP clones in an offline+FmiΔcad wing. N=10 individual wings from different animals G. Vang::YFP clone near the anterior crossvein in the region of the swirl. The location corresponds to the yellow box in Supplementary Figure 1B. H, I. Vang::YFP clones visualized with TIRF microscopy in wild-type Fmi or offline wings. N=10 individual wings from different animals for each condition. J. Polarization scores, defined as the ratio of averaged fluorescence intensity on the bright sides over the average intensity on the opposite sides of an individual clone (see Methods) for Vang::YFP and Fz::YPF clones in wildtype and offline wings. Scores greater than 1 (dashed line) represent polarized clones. Significance was tested with a one way ANOVA to compare mean variance. *** denotes p < .001; ** denotes p < .01. Center line represents the mean. Scale bars = 5 μm.

In single cells that polarize, theory suggests that symmetry breaking requires both positive and negative feedback^22,23^, and experimental evidence from shmoo formation in budding yeast, chemotaxis of neutrophils and *Dictyostelium*, T cell activation, and developmental cell migration supports this view^24–26^. In these systems, positive feedback acts to localize and reinforce the accumulation and activation of signaling molecules that mark a pole of the cell, while negative feedback excludes or suppresses the accumulation and activation of those factors in other locations. In neutrophils and *Dictyostelium*, two such interlocking systems localize to opposite poles and reinforce each other. For example, yeast cooperatively recruit and activate Cdc42, resulting in activation of Cdc42-GTP in a focused spot of signaling and reduction of signaling elsewhere^24^. Similarly, migrating neutrophils exhibit positive feedback of Rac activation, mediated by PIP3 and Cdc42, producing a leading protrusive edge^25^. In these systems, clustering of key signaling proteins is a critical driver of positive feedback.

In PCP, both positive and negative feedback may also underlie symmetry breaking and polarization^21,22^ (Fig. 1A-B). Positive feedback might arise via several mechanisms, such as cooperative assembly of PCP complexes into clusters or reinforcement of localization, or through double-negative loops in which the two subcomplexes antagonize each other (known as mutual inhibition). The presence of negative feedback is supported by indirect evidence deriving from mosaic analyses^14,27^. Because PCP is intrinsically multicellular, possibilities for positive and negative feedback exist that are not available to isolated polarizing cells. In PCP, the operative feedback pathways are not well defined, nor is it known whether the ability of PCP to polarize cells independent of intercellular communication is conserved.

Here, we demonstrate that in the *Drosophila* wing, cells deprived of their ability to communicate polarity information from cell to cell can nonetheless polarize. We provide evidence for one negative feedback mechanism and possible positive feedback mechanisms, and show that at least some of these function in the absence of intercellular polarity communication. Finally, we propose a mechanism for symmetry breaking involving negative feedback via competitive interactions.

## Results

### A tool to eliminate intercellular PCP signal transmission

Because Fmi-mediated intercellular PCP communication is a central feature of core PCP function^14–18^, it is not apparent whether cells incapable of communicating polarity information through Fmi can polarize. To address this, we developed a genetic tool that confers the ability to ‘isolate’ individual cells from their neighbors with respect to PCP information (here referred to as being ‘offline’). It is firmly established that Fmi homodimers are essential for communication between Van Gogh/Prickle and Frizzled/Dishevelled (proximal and distal, respectively, with respect to the wing) complexes on neighboring cells^14–18^. To block intercellular PCP communication, we remove Fmi entirely using *fmi* null alleles, or replace endogenous Fmi with a construct that disrupts *trans* homodimerization but should still facilitate intracellular complex assembly (FmiΔcad; lacking all cadherin repeats). To preserve viability^28^, we rescue CNS Fmi expression using the pan-neural GAL4-1407 converted to the Q system^29^. We verified absence of QF2-1407 activity in the wing by crossing to QUAS-GFP, and the absence of Fmi in the wing by antibody staining (Supp Fig. 1A). As anticipated, adult offline wings show a strong polarity phenotype with stereotypical swirling patterns and discontinuities similar to those of other core PCP mutants (Supp Fig. 1C-E). We thereby employ a truly tissue-wide null condition whereas hypomorphic alleles were the only option available for prior studies.

### Offline cells polarize

To determine whether individual offline cells polarize, Vang::YFP and Fz::YFP were mosaically expressed in wild-type and offline backgrounds, enabling us to unambiguously determine the distribution of Vang and Fz within individual cells. Remarkably, in offline cells in various regions of the wing, as in the wild-type background, individual cells showed clear asymmetric segregation of both Vang and Fz, albeit the signals were weak, perhaps in part due to dilution by unlabeled endogenous protein (Fig. 1C-E”). Furthermore, in the offline background the direction of cellular polarization was not coordinated across the tissue, consistent with the mutant adult hair polarity pattern. Pre-hairs emerged from the side of the cell opposite to Vang::YFP and the same side as Fz::YFP localization, as occurs in wildtype (Fig. 1D-E” and Supp Fig. 2). In the offline background, we then expressed FmiΔcad, which lacks the first 8 of 9 cadherin repeats and therefore cannot trans-homodimerize^30–32^, but should retain the ability to interact with other core components cell-autonomously. As expected, supplying FmiΔcad failed to correct the strong offline polarity defect (Supp Fig. 1E), and as in *fmi^null^* wings, cells polarized as judged by asymmetric Vang::YFP localization (Fig 1F).

Clustering of core PCP components is associated with polarization^27,33^, and in related work from our lab is shown to be required for polarization^34^. Since Fmi has been proposed to be a scaffold upon which PCP complexes form^17,18^, in its absence one might expect no clusters (a.k.a. puncta) to form and polarization to fail. Because we observed polarization, we examined whether PCP clusters form in the absence of Fmi. Some apparent clustering of Fz::YFP and Vang::YFP may be seen by confocal microscopy in the offline, or offline plus FmiΔcad wings (Fig. 1D-G), but the limited resolution and weak signal make this observation inconclusive. We therefore imaged the same offline wings using live Total Internal Reflection Fluorescence (TIRF) microscopy as described in a related manuscript^34^. We observed polarization similar to that seen with confocal microscopy, though clustering is more apparent (Fig. 1H-I). To quantify polarity of both Fz::YFP and Vang:YFP in offline wings, using the confocal images, we determined a polarization score, defined as the ratio of signal on the bright side over signal on the opposite side (see Methods); a value of 1 indicates no polarity (Fig. 1J). Indeed, our analysis demonstrated both Fz and Vang polarize in offline wings, albeit less well than their WT counterparts.

From these results, we conclude that communication of polarity information between cells is not required for cells to polarize, though their direction of polarization fails to follow the tissue axes. Furthermore, polarization is accompanied by clustering, at least of Vang, despite the absence of Fmi.

Mutations that disrupt core PCP signaling do not produce randomized polarity of the wing hair pattern despite eliminating detectable asymmetry of core protein localization. Rather, characteristic regions of whorls and more uniform polarity are seen in core PCP mutants. The determinants of this pattern are not well understood, but evidence points to mechanical factors and is described in the Discussion. The wing hair pattern of offline wings strongly resembles that of other core mutants (Supp Fig. 1D), though unlike other core mutants, in offline wings asymmetric localization of core proteins occurs, and their orientation correlates to the orientation of hairs. In regions of offline wings where polarity is relatively uniform, so too is the direction of Vang::YFP polarization. In contrast, in regions where the hair polarity is most chaotic, asymmetry of Vang::YFP is difficult to discern (Fig. 1G). From these observations, we infer that the direction of offline core protein polarization is likely determined by the same, possibly mechanical, cues that determine the mutant hair polarity pattern.

### Feedback in PCP signaling

In biochemical analyses of Wnt signaling, Dvl2 (vertebrate Dsh) recruitment by Fz depends on positive feedback involving PI (4,5)P2^35^. The potential existence of a similar positive feedback mechanism in PCP signaling has not been evaluated. We hypothesized that positive feedback might occur by mutually reinforcing recruitment, clustering and/or stabilization between Fz and Dsh. During PCP signaling, Fz is required to recruit Dsh to the plasma membrane^36–38^, and overexpression of Pk causes excess recruitment of Dsh, Fz and Fmi, also consistent with positive feedback^39^. We therefore wished to test for the possibility of a self-promoting loop between Fz and Dsh by determining whether Fz recruitment, stabilization or clustering might depend on Dsh.

Evidence for negative feedback in PCP signaling derives from mosaic experiments in which clonal overexpression of Vang, which induces recruitment of Fz on neighboring cell boundaries, excludes Vang from the neighboring cell boundaries^14^; exclusion of Vang requires the presence of Pk^27^. In a converse experiment, we here show that clonal overexpression of Fz recruits Vang to the neighboring cell boundary and excludes Fz^14^, and this also requires the presence of Pk (Supp Fig. 3A-D’’). Both of these experiments rely on intercellular signaling, and while it was hypothesized that the mutual exclusion between Fz and Vang might occur cell-autonomously, the presence of intercellular signaling allows that it may have occurred through an indirect route. Therefore, the direct interactions that lead to mutual exclusion remain to be defined^21,40,41^ (red arrows in Fig. 1A).

To explore more direct relationships between PCP protein localization and effects on other PCP components, particularly those occurring within the same cell, we developed a genetic ‘Velcro’ approach to enforce a local asymmetry within the core PCP complex even when PCP signaling is otherwise disrupted. We then assay which other core proteins are either recruited or excluded by the localized protein.

The Velcro assay uses N-Cadherin fused to the transmembrane and intracellular domains of Fz (N-Cad::V5-Fz), or to an anti-myc scFv^42^ (N-Cad::HA-3DX) which recruits any myc-tagged protein to cell-cell junctions (Fig. 2A,B). Expressing N-Cad::V5-Fz or N-Cad::HA-3DX in clonal cell populations in the pupal wing recruits N-Cad::V5-Fz or Dsh-myc selectively to cell-cell boundaries where the neighboring cell also expresses the N-Cad construct, thereby enforcing asymmetry in cells at the periphery of the clone (Fig. 2C,D). In the Fz Velcro wings, the N-Cad::V5-Fz is the only Fz in the tissue, such that the background is unpolarized. In contrast, Dsh-myc is uniformly expressed and supports normal polarization where not perturbed by N-Cad::HA-3DX clones. As expected, hair polarity is reoriented by both Velcroed Fz and Velcroed Dsh (Supp Fig. 4A-D).

**Figure 2.**
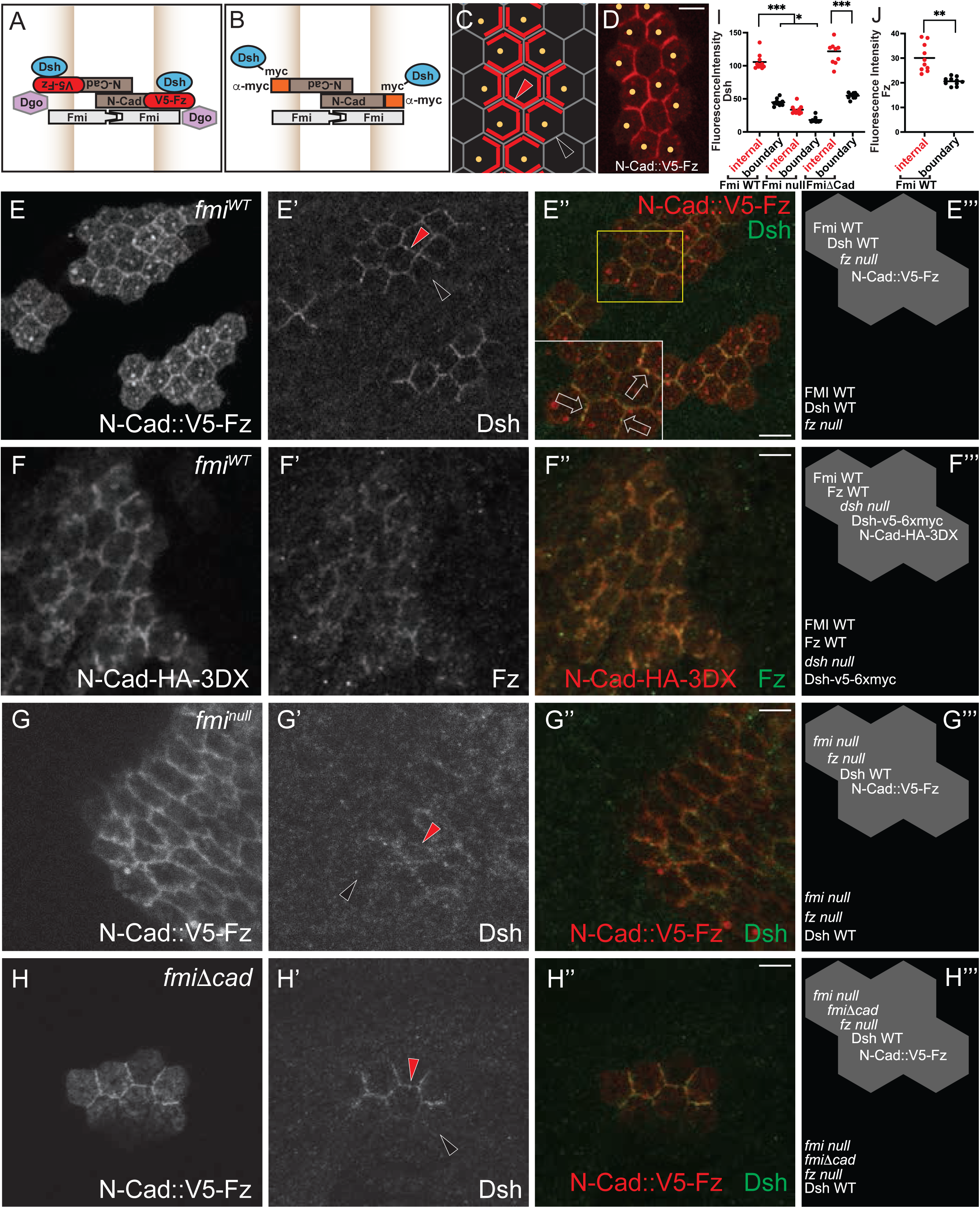
Velcroed Fz recruits Dsh and Velcroed Dsh recruits Fz in wildtype and offline backgrounds. A. Schematic of the Fz Velcro system. Clonal N-Cad::V5-Fz expression is expected to recruit N-Cad::V5-Fz selectively to shared cell junctions (C,D). No endogenous Fz is present. B. Schematic of the Dsh Velcro system. Clonal expression of N-Cad::HA-3DX in wings uniformly expressing Dsh-myc at low levels is expected to recruit Dsh-myc to shared cell junctions. C. Schematic of clonal expression (yellow dots outline hypothetical clone) recruiting Velcroed components (red) to shared cell junctions. At the periphery of the clone, recruitment is expected to be to some but not all cell junctions D. N-Cad::V5-Fz stained with anti-V5, showing localization to shared cell junctions. E-E’’’. N-Cad::V5-Fz clones in a *fz^null^* background (Velcroed Fz) stained for V5, Dsh (E’) and merged (E’’). N=10 individual wings from different animals Note that the Dsh antibody is not sensitive enough to detect endogenous Dsh, but little is expected in the surrounding tissue because it is mutant for *f*z. F-F’’. N-Cad::HA-3DX clone in a Dsh-myc expressing wing (Velcroed Dsh) stained for HA to detect N-Cad::HA-3DX (F), Fz (F’) and merged F’’. N=10 individual wings from different animals Note that the Fz antibody is not sensitive enough to detect endogenous Fz, so it only detects Fz colocalizing with N-Cad::HA-3DX where it is recruited to sufficient levels for detection. G-G’’. N-Cad::V5-Fz clones in *fz^null^*, *fmi^null^* background (Velcroed Fz, offline) stained for V5 (G), Dsh (G’) and merged (G’’). H-H’’. N-Cad::V5-Fz clones in *fz^null^*, *fmi^null^*, arm-fmiΔcad background (Velcroed Fz, offline with FmiΔcad) stained for V5 (H), Dsh (H’) and merged (H’’). N=10 individual wings from different animals E’’’-H’’’. Cartoons of genotypes inside and outside the clones in E-H. I. Quantification of the cell boundary intensity linescans internal to the clone and at clone boundaries as in C, E’, G’, H’. Dots are color coded to the arrowheads in showing examples of each (see Methods). J. Quantification of the cell boundary intensity linescans internal to the clone and at clone boundaries as in F’. Significance was tested with a one way ANOVA to compare mean variance. *** denotes p < .001; ** denotes p < .01. * Fmi WT boundary compared with Fmi Null Boundary p = .034. Center line represents the mean. Scale bars = 5 μm.

This method allowed us to assay several potential feedback mechanisms that depend on interactions within one cell. We first examined the potential recruitment of Dsh by Fz, as suggested by prior results^36,38^, and *vice versa*, as is expected if positive feedback is occurring cell-autonomously. To do so, we Velcroed Fz and assayed for Dsh recruitment. To monitor Dsh, we used an antibody that has sufficient sensitivity for this purpose^43^. N-Cad::V5-Fz was strongly expressed under UAS control, and where N-Cad::V5-Fz localized, we observed recruitment of endogenous Dsh (Fig. 2E-E’’’,I). When N-Cad::V5-Fz is simultaneously monitored, substantial co-localization of N-Cad::V5-Fz and Dsh are observed in puncta (Fig. 2E”). We then did the reciprocal experiment, Velcroing Dsh-myc and monitoring Fz. The Fz antibody does not detect endogenous levels of Fz, but reliably, though only weakly, detected elevated levels of Fz where Dsh-myc is localized (Fig. 2F-F”’,J). The Dsh Velcro system is weaker than the Fz Velcro system, in part because Dsh-myc is expressed at low levels, and in part because endogenous Myc competes with Dsh-myc for binding to N-Cad::HA-3DX (Supp Fig. 4E,E’). Recruitment of Dsh by Velcroed Fz, and recruitment of Fz by Velcroed Dsh suggest the possibility of positive feedback occurring via this loop.

In addition to assaying mutual recruitment of Fz and Dsh, the Velcro assay allowed us to assay a potential mechanism for negative feedback: exclusion of the Vang/Pk complex from the locations where N-Cad::V5-Fz or Dsh-myc is localized (“internal” boundaries Fig. 3A). Such exclusion could be inferred from the known opposing localizations of the Vang/Pk complex and the Fz/Dsh complex, and from more elaborate assays (Supp Fig. 3 and ref ^27^), but prior assays could not distinguish whether exclusion requires information from the neighboring cell or if localized Fz without a signal from the neighbor is sufficient to mediate exclusion. For these assays, we chose the stronger N-Cad::V5-Fz Velcro, and assayed Pk rather than Vang due to the availability of a much more sensitive antibody. Three possible outcomes were envisioned: *i)* no exclusion, in which case Pk localizes uniformly around the cell (black arrow; Fig. 3B); *ii)* exclusion results in only cytoplasmic signal (negative feedback; green arrow); and *iii)* exclusion from Fz-occupied boundaries and ‘capture’ at unoccupied boundaries (yellow arrow).

**Figure 3.**
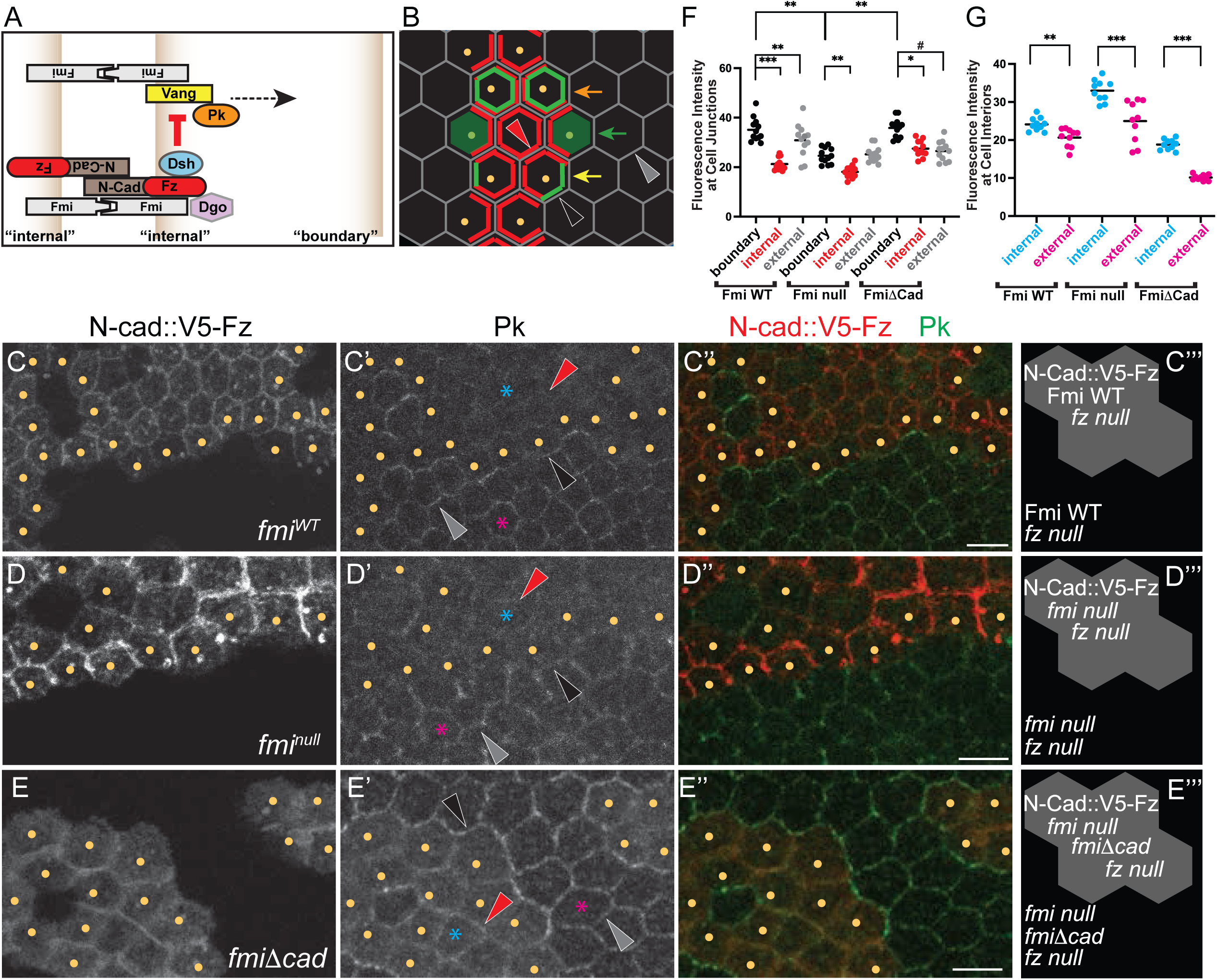
Velcroed Fz excludes Pk in wildtype and offline backgrounds. A. Schematic of Velcroed Fz excluding the Vang-Pk subcomplex. B. Potential outcomes of the exclusion experiment showing localization of Pk: no exclusion (orange arrow), exclusion and accumulation in cytoplasm (green arrow), and exclusion and capture of Pk at cell boundaries lacking N-Cad::V5-Fz (yellow arrow; clone “boundary”). To quantify Pk capture at clone boundaries, intensity line scans were done at clone boundaries (white arrowhead) and compared to intensity linescans of nearby external cell boundaries (gray arrowhead). C-C’’’. N-Cad::V5-Fz clones in *fz^null^* background (Velcroed Fz) stained for V5 (C), Pk (C’) and merge (C’’). N=10 individual wings from different animals. Pk is excluded from internal clone boundaries where N-Cad::V5-Fz localizes (red triangle) and cytoplasmic level of Pk is increased. Pk is captured at clone boundaries (quantified in F; because Pk from the cell outside the clone also may accumulate at that cell interface, a contribution due to exclusion from the clone cell was scored as a level higher than at external cell junctions). D-D’’’. N-Cad::V5-Fz clones in *fz^null^*, *fmi^null^* background (Velcroed Fz, offline) stained for V5 (D), Pk (D’) and merge (D’’). N=10 individual wings from different animals. Pk is excluded from internal clone boundaries where N-Cad::V5-Fz localizes and cytoplasmic level is increased. Pk is excluded from internal clone boundaries where N-Cad::V5-Fz localizes and cytoplasmic level is increased. Pk is not captured at clone boundaries (quantified in F). E-E’’’. N-Cad::V5-Fz clones in *fz^null^*, *fmi^null^*, arm-fmiΔcad background (Velcroed Fz, offline with FmiΔcad) stained for V5 (E), Pk (E’) and merge (E’’). N=10 individual wings from different animals. Pk is excluded from internal clone boundaries where N-Cad::V5-Fz localizes and cytoplasmic level is increased. Pk is excluded from internal clone boundaries where N-Cad::V5-Fz localizes and cytoplasmic level is increased. Pk is captured at clone boundaries (quantified in F). C’’’-E’’’. Cartoons of genotypes inside and outside the clones in C-E. F. Quantification of cell boundary intensity linescans as described in B for the results in C-E (see Methods). Dots are color coded to the arrowheads in showing examples of each (see Methods). G. Quantification of cell interior areas in C’, D’, E’. Asterisks are color coded to the arrowheads in showing examples of each (see Methods). Significance was tested with a one way ANOVA to compare mean variance. *** denotes p < .001; ** denotes p < .01. *fmi^null^*, arm-fmiΔcad boundary compared with internal (*) and external (#) were p = .035 and p = .022, respectively. Center line represents the mean. Scale bars = 5 μm.

Indeed, in cells where Fz localization was enforced at all cell edges, cytoplasmic Pk was elevated compared to cells outside the clone (Fig. 3C-C’’,G), and little Pk was observed at the plasma membrane (Fig. 3C-C”,F); red arrowhead). In cells where Fz localization was forced to only some cell edges (i.e. at the clone periphery), Pk was not only excluded from N-Cad::V5-Fz occupied edges, but localized selectively to outside cell edges where N-Cad::V5-Fz is not localized (Fig. 3C; black arrowhead and Fig. 3F). These observations indicate that enforced localization of Fz that is not expected to be interpreting polarity information from the neighboring cell is capable of excluding Pk. Together with this negative feedback, the reciprocal, direct or indirect negative feedback embodied by Vang’s exclusion of Fz described above (Supp Fig. 3D-D’’) form a double negative, and therefore positive, feedback loop.

### Feedback in offline cells

Because offline cells polarize (Fig. 1D-E), we assayed whether the signs of positive and negative feedback just described might also occur in offline conditions by deploying the N-Cad::V5-Fz Velcro system in offline wings. Assays testing whether these feedback mechanisms operate cell autonomously have not previously been possible. In the absence of Fmi, N-Cad::V5-Fz recruits Dsh, though less efficiently than in the presence of Fmi (Fig. 2G-G”’,I). In the presence of FmiΔcad, Dsh was recruited nearly as efficiently as in the presence of intact Fmi (Fig. 2H-I). Therefore, consistent with the inference that positive feedback must occur in offline cells, we see one facet of positive feedback in the cell autonomous recruitment of Dsh by Fz.

Similarly, exclusion of Pk by N-Cad::V5-Fz occurs in offline cells and therefore occurs cell autonomously. N-Cad::V5-Fz in the absence of Fmi increases the level of Pk that resides in the cytoplasm, and little Pk remains at cell boundaries occupied by N-Cad::V5-Fz (Fig. 3D-D’’,F,G). N-Cad::V5-Fz in the presence of FmiΔcad also causes additional Pk to remain in the cytoplasm (Fig. 3E-E’’’,G). Exclusion therefore occurs independent of cell-cell PCP communication. Of note, without Fmi, the excluded Pk is not appreciably captured at the unoccupied outside boundaries, while presence of FmiΔcad facilitates its capture (Fig. 3D’; black arrowhead, and Fig. 3F). We conclude from these experiments that negative feedback embodied in the exclusion of Pk by Fz occurs in the absence of cell-cell PCP communication.

### Negative feedback requires Dsh-dependent clustering

Since asymmetry is thought to arise in PCP puncta, we asked whether Velcroed Fz-mediated exclusion of Pk requires clustering. To do so, we employed a rationally designed Dsh mutant, Dsh^G63D^. The orthologous mutation in the mouse Dishevelled2 (Dvl2) DIX domain impairs oligomerization by disrupting antiparallel inter-strand interactions^44^, and *dsh^G63D^* is shown in a related manuscript to allow clusters to form but to decrease the size of clusters, and to produce a strong polarity phenotype^34^. We find that Velcroed Fz in the *dsh^G63D^* mutant background fails to exclude Pk (Fig. 4A-B”’,E, and Supp Fig. 5), suggesting that Dsh-dependent assembly of large clusters is required for negative feedback via Pk exclusion.

**Figure 4.**
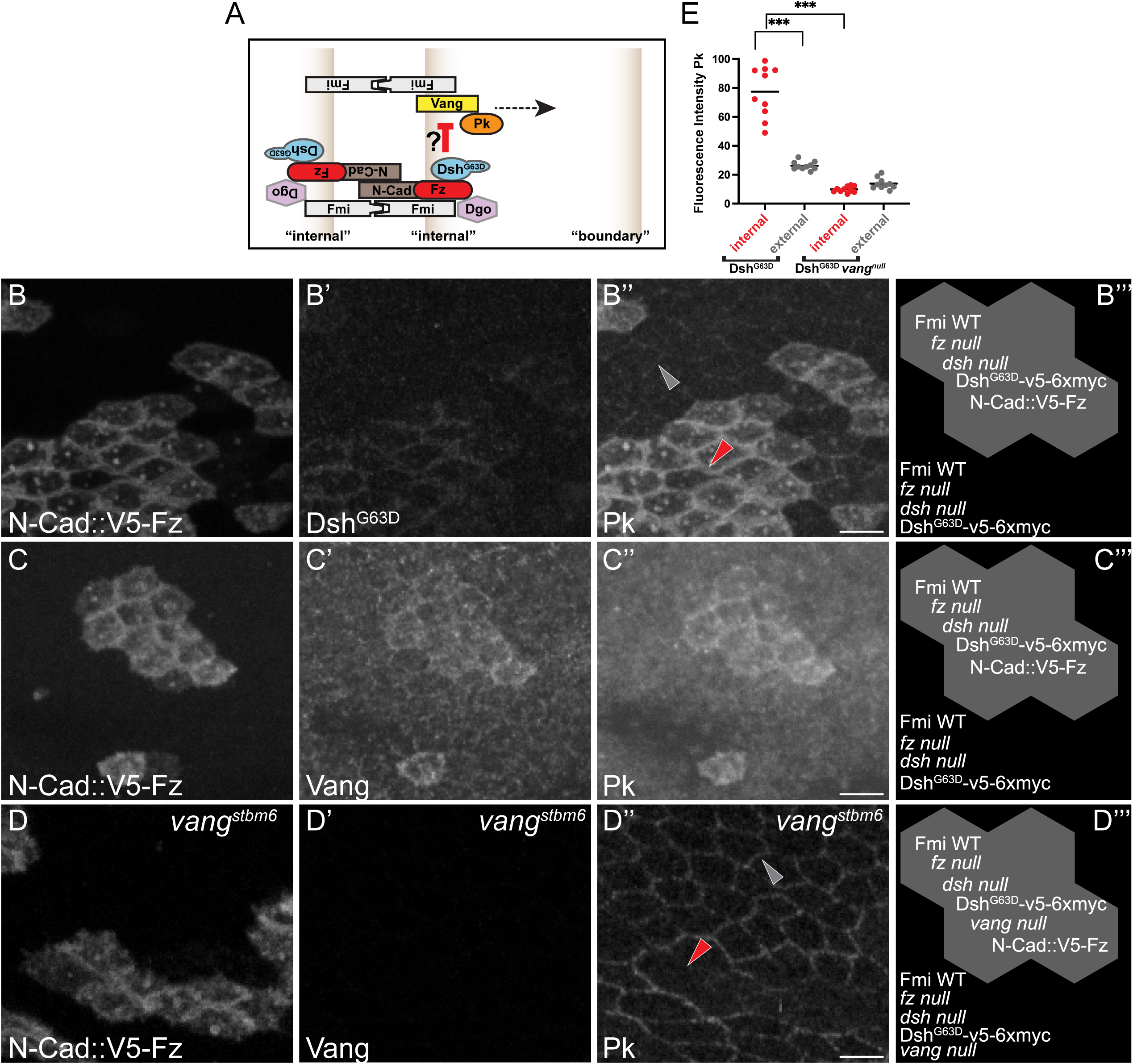
Negative feedback requires Dsh-dependent clustering. A. Schematic of Velcroed Fz failing to exclude the Vang-Pk subcomplex in a Dsh^G63D^ background. B-B’’. N-Cad::V5-Fz clones in a *dsh^G63D^*, *fz^R^*^52^ mutant background, stained for V5 (N-Cad::V5-Fz; B), Dsh (B’) and Pk (B’’). N=10 individual wings from different animals. Dsh^G63D^ is recruited to Velcroed Fz, but in the presence of a clustering deficient Dsh, fails to exclude Pk. Furthermore, Pk is recruited to higher levels than in the *fz* mutant background. For an essential control, see Supp Fig. 4. C-C’’. N-Cad::V5-Fz clones in a *dsh^G63D^*, *fz^R^*^52^ mutant background, stained for V5 (N-Cad::V5-Fz; C), Vang (C’) and Pk (C’’). N=10 individual wings from different animals. Vang is recruited to Velcroed Fz, perhaps mimicking the population of distal Vang that is observed in wild type cells. D-D’’. N-Cad::V5-Fz clones in a *dsh^G63D^*, *fz^R^*^52^, *vang^stbm^* mutant background, stained for V5 (N-Cad::V5-Fz; D), Vang (D’) and Pk (D’’). N=10 individual wings from different animals. In the absence of Vang, Velcroed Fz with a clustering deficient Dsh no longer recruits Pk. B’’’-D’’’. Cartoons of genotypes inside and outside the clones in B-D. E. Quantification of the cell boundary intensity linescans internal to the clone and at clone boundaries as in B-D. Dots are color coded to the arrowheads in showing examples of each (see Methods). Significance was tested with a one way ANOVA to compare mean variance. *** denotes p < .001. Center line represents the mean. Scale bars = 5 μm.

For reasons not readily explained by current models, not only is Pk not excluded by Velcroed Fz, but Pk is recruited to higher levels than in the surrounding cells by Velcroed Fz in a *dsh^G63D^*mutant background (Fig. 4C-C’’’ and Supp Fig. 5). To further understand this result, we considered another surprising observation from a related manuscript from our lab showing that a substantial population of Vang is present in clusters on the distal side of the cell where Fz accumulates^34^. We hypothesized that if Dsh-dependent clustering is required for Fz to exclude Pk, the recruited Pk in the *dsh^G63D^* mutant might result from the presence of a ‘distal’ Vang population alongside the Velcroed Fz. Indeed, a population of Vang is present together with Velcroed Fz (Fig. 4C-C’”), and Velcroed Fz in the *dsh^G63D^* mutant does not recruit Pk if Vang is also removed (Fig. 4D-E). This suggests that in at least some endogenous PCP clusters, a population of distal Vang is present alongside Fz and is capable of recruiting Pk, but that normally, Fz recruits Dsh that both promotes PCP clustering and excludes Pk. We propose below a model for how this may contribute to symmetry breaking within puncta.

## Discussion

Fmi homodimers bridging between cells are thought to be the scaffold upon which core PCP complexes are assembled^14,17,18,45^. A central role for Fmi homodimers in scaffolding PCP complexes and in communicating PCP information between cells might suggest that the positive and negative feedback leading to polarization is inextricably linked to intercellular communication^14–18^. Recent work by Basta et. al. 2025, in a chimeric mouse epidermal skin model, argues just this, that PCP polarization is not cell-autonomous, and requires intercellular coordination through Celsr1/Fmi. Yet, we show here that in *Drosophila* wings, individual cells polarize in the absence of Fmi. The reason for the difference between our results and those of Basta et al is not immediately apparent; while it is possible that PCP signaling is fundamentally different in the two systems, there are other potential explanations worth entertaining. First, it is possible that the difference is quantitative, such that any autonomous polarization of mouse epithelial cells is below their threshold of detection. Second, different approaches could account for a difference in outcome, as Basta et. al. surrounded cells expressing Celsr1 with cells mutant for Celsr1, whereas our most comparable condition replaced Fmi with the communication deficient FmiΔCad. At a conceptual level, however, it is hard to imagine why this difference should be consequential, as one would expect CELSR without a partner in the neighboring cell to support functionality in a manner equivalent to FmiΔCad. Based on our results, we conclude that sufficient feedback operates within individual cells to cause them to polarize autonomously. The logic of the PCP system thus may more closely resemble that of single cell polarizing mechanisms in yeast, neutrophils, *Dictyostelium*, T cells and *C. elegans* embryos^24–26,46^ than might otherwise have been expected.

We provide evidence for both negative and positive feedback in core PCP polarization, and argue that at least some of this feedback can operate in ‘offline’ cells. Our data demonstrate negative feedback in the exclusion of the proximal complex component Pk from domains where localization of the distal domain component Fz is enforced. Exclusion of Pk by Fz occurs cell-autonomously, and likely depends on clustering, since it depends on the ability of Dsh to oligomerize and promote clustering (Figs. 3 and 4). Vang is also cell-autonomously excluded by localized Fz, perhaps by removal of Pk, which may destabilize Vang, though this is apparently less complete than the exclusion of Pk (Supp Fig. 6A-C). Conversely, negative feedback is also apparent in the previously demonstrated exclusion of Fz from domains where Vang is enriched^14,27^; currently available tools do not allow us to assay whether Fz exclusion occurs cell-autonomously. Exclusion of Fz by Vang and *vice versa* each depend on the presence of Pk (Supp Fig. 2 and ref.^27^), and we suggest this involves a mechanism in which Pk and Dsh compete for Vang binding.

We suggest two potential positive feedback mechanisms. First, the two negative feedbacks embodied in the mutual exclusion of proximal and distal components together form a loop that will produce positive feedback. Second, the enhancement of Dsh recruitment by Fz, and Fz by Dsh, into clusters might also produce positive feedback.

Intercellular Fmi bridges transmit an asymmetric signal in which one cell tells its neighbor that Fz is present at its surface, thereby recruiting Vang to the neighbor, and at the same time, the Vang in the neighbor cell tells the first cell that Vang is on its surface, eliciting recruitment of Fz to its surface^14,47^. Fmi, by virtue of it being a cadherin, also provides an adhesive interaction between the opposite complexes. Our argument that the feedback mechanisms we examined can function in the absence of polarity information from neighboring cells depends on the use of Velcroed Fz. In the Velcro scenario, N-Cad::Fz localizes Fz using the adhesive function of N-Cad; we argue that this adhesive function cannot replace the asymmetric signaling through Fmi that is eliminated when Fmi is removed (offline). N-Cad::Fz is present on both sides, so there is no potential for any asymmetric signaling. Despite the presence of N-Cad::Fz in the neighbor, N-Cad::Fz behaves like Fz, recruiting Dsh and excluding Pk, a response that would not be expected if it were responding to Fz in the neighboring cell. A further, though more subtle, argument can also be made. One might be tempted to argue that the Vang we detect in the N-Cad::Fz complexes is signaling through N-Cad in trans to induce the observed behavior of N-Cad::Fz, but we know that this cannot be true because Pk is still excluded when there is no Vang present (Fig 4D). We are therefore confident that N-Cad::Fz localizes N-Cad::Fz in the neighbor but provides no signaling information. The feedback we characterize in the Velcro system is thus likely to be operative in offline cells.

We do not mean to imply that the feedback loops identified here are exclusive of other potential feedback loops. For example, it is possible that positive feedback promotes assembly of the Vang-Pk complex, and additional feedback pathways involving intercellular communication are likely to operate. Furthermore, it is likely that some form of long-range negative feedback is necessary. Redundancy in feedback pathways would contribute to the reliable function of PCP polarization and the stabilization of clusters. Regardless of the full complement of feedback that underlies PCP signaling, if theory holds^22,23^, one can conclude that in offline cells, at least some positive and negative feedback occurs.

Because Fmi was thought to template the assembly of PCP signaling complexes^14,17,18,45^, the ability of cells to polarize in the absence of Fmi was unexpected. This implies that the other core proteins retain some ability to interact independently of Fmi. Many physical interactions between core proteins have been documented by co-immunoprecipitation that are consistent with this possibility. Fmi likely provides a helpful, albeit non-essential, scaffolding function to facilitate polarization of individual cells.

As noted above, strong mutations of core polarity components produce very similar whorl patterns of adult wing hairs, and this observation now extends to the *fmi^null^* (offline) condition. While the determinants of this pattern are not well understood, experiments that modify the mechanical environment by eliminating wing veins substantially simplify the hair polarity pattern of PCP mutants^48^. Because vein cells have smaller apical surfaces than other cells, the cell packing patterns are necessarily less regular near veins, and additional evidence has shown that cell packing contributes to polarization and *vice versa*^49,50^. Thus, the core mutant wing hair polarity pattern may derive from the cell packing pattern. Additional mechanical influences across the developing wing, such as cell flow have also been proposed^51^. In vertebrate systems, tissue deformation via tension has also been shown to orient PCP patterns^52–54^. In one case, this was shown to depend on microtubules^54^. In the fly wing, since microtubules both contribute and respond to cell shape^55^, and apical microtubules are proposed to provide directional bias to core PCP polarization^56^, microtubules may have a role in this process. Notably, despite cells polarizing their core proteins in offline cells, the aberrant hair polarity pattern remains uncorrected. We propose that offline cells neither align their polarities via Fmi-mediated intercellular interactions nor respond to global directional signals.

In a related manuscript, we observed a population of distal Vang in endogenous PCP complexes^34^, and the presence of Vang together with Velcroed Fz suggests that Velcroed Fz provides a model that mimics distal Vang. We propose that as a complex is first forming, and as it subsequently evolves, through a stochastic process both Fz or Vang can partner with Fmi on either side. A mechanism is then required during symmetry breaking to attenuate the activity of Vang that may have been incorporated on the distal side. Because, in the wing, Fz and Dsh are delivered in a biased fashion to the distal side^57–60^, in time, Fz is likely to enter the complex, subsequently recruiting Dsh. Our exclusion results indicate that the presence of Dsh excludes Pk from these complexes, likely through competitive binding to Vang, and in a fashion requiring its ability to oligomerize^34^. We suggest that excluding Pk antagonizes Vang functionality and allows Fz to fortify the distal character of the complex. Dsh and Pk have been shown to competitively bind Vang in a phosphotyrosine dependent fashion^45,61,62^. The relationship between tyrosine phosphorylation, Fz recruitment of Dsh, and Dsh competing with Pk for Vang binding remain to be determined. Once established, distal Fz facilitates cluster growth, including accumulation of more proximal Vang within that cluster^34^.

Whereas the wing depends on the Pk^pk^ isoform, we speculate that in Pk^sple^-dependent tissues, the competition between Pk and Dsh for Vang binding is influenced in a different fashion. Pk^sple^ is known to bind and be recruited to the side of the cell enriched for Dachsous and Dachs^63,64^, and we suggest that this recruited Pk then out-competes Dsh for binding to Vang, thereby promoting the Fmi-Vang-Pk complex on that side of the cell.

While we demonstrate exclusion of Pk by Fz, the present results provide no clear evidence for or against Fz excluding Vang. Indeed, current evidence suggests a more complex relationship. Clonal Vang overexpression, which recruits Fz to the adjacent wildtype cell boundary, results in exclusion of most Vang from that boundary, and its exclusion requires Pk^27^, so it is reasonable to infer that recruited Fz has at least some capacity to exclude Vang. On the other hand, Velcroed Fz is seen to associate with some Vang in cis (Fig. 4B’), yet appears to partially exclude Vang as well (Supp Fig. 6). In a related manuscript, it is shown that Vang is measured together with Fz in at least some clusters in modest amounts that do not correlate to the amount of other components^34^. These observations taken together suggest that while there may be some affinity for Vang in distal clusters, as clusters grow, distal Vang tends to be excluded. Additional work will be required to better understand the regulation of Vang in distal complexes.

In *Drosophila*, evidence of a physical interaction between the Fz extracellular domain and Vang was used to propose a trans interaction^65^; based on results here and in a related manuscript^34^, we suggest instead that they interact in cis. Fz, Vang, Dsh and Pk orthologs have been proposed to collaborate in cis in at least one other context as well. In Xenopus animal cap and dorsal marginal zone cells undergoing convergent extension, a competitive, signal dependent competition has been proposed to regulate the association of Dvl2 with either Vangl2 or Fz7^66,67^.

We show here that the *Drosophila* PCP system retains the ability to polarize cells cell-autonomously when stripped of its ability to transmit polarity information between neighboring cells. Negative and potential positive feedback mechanisms are identified that could support this polarization, and our results suggest a mechanism for establishing and maintaining distal character by competitive binding of Dsh and Pk to Vang. Future studies will be required to further characterize these feedback mechanisms and to define other intracellular and intercellular mechanisms that are likely to also contribute to polarization.

## Methods

### Drosophila Husbandry and Genetics

Flies stocks were raised on standard fly food at 25°C. Heat shock protocols were performed on third instar larvae at 37°C and moved back to 25°C until selection of pupae. Standard *Drosophila* genetic techniques were used to create recombinant chromosomes. Compound balanced fly stocks were then generated to create stable fly lines.

Homology assisted CRISPR knock-in (HACK) was used as described in Lin and Potter^29^ to convert 1407-GAL4 to QF2-1407. QF2-1407 was validated by driving QUAS-eGFP; we observed the expected CNS expression pattern, but no signal in larval or pupal wings.

### Drosophila genotypes

Genotypes of experiments by figure are listed below.

Figure 1:

Figure 1C and 1H:

+/+; hsFLP; TM2, act>>vang::YFP

Figure 1C:

+/+; hsFLP; act>>fz::YFP

+/+; hsFLP; +/TM2, act>>vang::YFP

Figure 1D-D”, G and I:

*yw* hsFLP; QF2-1407, *fmi^E^*^59^ / *fmi^E^*^45^; QUAS-fmi (F17) / TM2, act>>vang::YFP

Figure 1E-E’’:

*yw* hsFLP; QF2-1407, *fmi^E5^*^9^ / *fmi^E4^*^5^; act>>fz::YFP, QUAS-fmi (F17)

Figure 1F:

*yw* hsFLP; QF2-1407, *fmi^E5^*^9^ / arm-fmiΔcad, *fmi^E4^*^5^; QUAS-fmi (F17) / TM2, act>>vang::YFP

Figure 1H:

+/+; hsFLP; +/TM2, act>>vang::YFP

Figure 2:

Figure 2D:

hsFLP; UAS-N-Cad::V5-Fz / act>>GAL4

Figure 2E-E”:

hsFLP; UAS-N-Cad::V5-Fz, *fz^R5^*^2^ / act>>GAL4, *fz^R^*^52^

Figure 2F-F”:

*dshV26*, dsh::V5-6xmyc; hsFLP; act>>GAL4 / UAS-N-Cad::HA-3DX

Figure 2G-G”:

*yw* hsFLP; QF2-1407, *fmi^E^*^59^ / *fmi^E^*^45^;

QUAS-fmi (F17), *fz^R^*^52^ / act>>GAL4, UAS-N-Cad::V5-Fz, *fz^R^*^52^

Figure 2H-H”:

*yw* hsFLP; QF2-1407, *fmi^E^*^59^ / arm-fmiΔcad, *fmi^E^*^45^;

QUAS-fmi (F17), *fz^R^*^52^ / act>>GAL4, UAS-N-Cad-V5-fz, *fz^R^*^52^

Figure 3:

Figure 3C-C”:

hsFLP; UAS-N-Cad::V5-Fz, *fz^R^*^52^ / act>>GAL4, *fz^R^*^52^

Figure 3D-D”:

*yw* hsFLP; QF2-1407, *fmi^E^*^59^ / *fmi^E^*^45^;

QUAS-fmi (F17), *fz^R^*^52^ / act>>GAL4, UAS-N-Cad::V5-Fz, *fz^R^*^52^

Figure 3E-E”:

*yw* hsFLP; QF2-1407, *fmi^E^*^59^ / arm-fmiΔcad, *fmi^E^*^45^;

QUAS-fmi (F17), *fz^R^*^52^ / act>>GAL4, UAS-N-Cad-V5-fz, *fz^R^*^52^

Figure 4:

Figure 4B-B” and 4C-C”:

Dsh^G63D^, *dsh^V^*^26^;; Act>>GAL4, UAS-N-Cad::V5-Fz, *fz^R^*^52^ / *fz^R^*^52^

Figure 4D-D”:

DshG63D, *dsh^V^*^26^; hsFLP, *stbm*^6^ / *stbm*^6^; Act>>GAL4, UAS-N-Cad::V5-Fz, *fz^R^*^52^ / *fz^R^*^52^

Supplementary Figure 1:

Supplementary Fig 1A-A” and D:

QF2-1407, *fmi^E^*^59^ / *fmi^E^*^45^; QUAS-fmi (F17)

Supplementary Fig 1C:

W118

Supplementary Fig 1E:

QF2-1407, *fmi^E^*^59^ / arm-fmiΔcad, *fmi^E^*^45^; QUAS-fmi (F17)

Supplementary Figure 2:

Supplementary Fig 2 A-A’’’

+/+; hsFLP; act>>fz::YFP

Supplementary Fig 2 B-B’’’

+/+; hsFLP; +/TM2, act>>vang::YFP

Supplementary Figure 3:

Supplementary Fig 3D-D”:

*hsflp/D174GAL4; FRT42D*, Fz::GFP, *tubP-GAL80 /FRT42D*, *pk ^pk-sple^*^13^*; UAS-mCherry/UAS-Fz*

Supplementary Figure 4:

Supplementary Fig 4C-C’ and 4F-F’:

hsFLP; UAS-N-Cad::V5-Fz, *fz^R^*^52^ / act>>GAL4, *fz^R^*^52^

Supplementary Fig 4D and 4F-F’:

hsFLP; *dsh^V^*^26^, dsh-myc; act>>GAL4, UAS-RFP / UAS-N-Cad::HA-3DX

Supplementary Figure 5

*Dsh^G63D^*, *Dsh^V^*^26^;; Act>>GAL4, UAS-N-Cad::V5-Fz, *fz^R^*^52^ / *fz^R^*^52^

Supplementary Figure 6

Supplementary Fig 6A and 6B:

*yw* hsFLP; QF2-1407, *fmi^E^*^59^ / *fmi^E^*^45^;

QUAS-fmi (F17), *fz^R^*^52^ / act>>GAL4, UAS-N-Cad::V5-Fz, *fz^R^*^52^

### Cloning and novel *Drosophila* strain construction

#### pQUAST-fmi

Full length *fmi* cDNA from UAS-fmi (ref ^5^) was subcloned in two EcoR1-Xho1fragments into pQUAST (#24349; Addgene) and transformed into *w^11^*^18^ flies to generate random genomic insertions.

#### pUAST-N-Cad::HA-3DX

A fragment of N-Cadherin cDNA (gift of Tom Clandinin) from the first aa to the SacII site, removing the C-terminus including the Δ-catenin binding site, was subcloned into pUAST. After mutating the internal SacII site, the HA-3DX sequence was generated by PCR (from ref ^42^) and appended to the SacII site in N-Cadherin. The resulting plasmid was transformed into *w^11^*^18^ flies to generate random genomic insertions.

#### pUAST-N-Cad::V5-Fz

The first 2915aa extracellular domain of N-Cadherin was generated by PCR from pUAST-N-Cadherin::HA-3DX using the following two primers:

Forward: AAGAGAACTCTGAATAGGGAATTGGGAATTCATGGCGGCACGACGTTGC

Reverse: CCGCCAGAGCCTGTGCCGCTATCACGGGATGGCGTCAATACAC

The resulting fragment contained the extracellular portion of N-Cadherin (2915aa) with the described linker and a V5 epitope tag.

The Fz transmembrane and C-terminus (final 347aa) was generated by PCR from (ref. ^68^) using the following primers:

Forward: ATCCCGTGATAGCGGCACAGGCTCTGGCGGCAAACCGATTCCGAACCCGCTGCTGGGCCTGGATAGCACCAGCGGAGGCGGTGGCATGTTCTTCCCGGAGAGAGAAAG

Reverse: CTTCACAAAGATCCTCTAGACTACACGTACGCCTGCGCCCG

The N-Cadherin and Fz fragments were cloned into EcoR1 and Xba1 digested pUAST.

The resulting plasmid was transformed into *w^11^*^18^ flies to generate random genomic insertions.

#### pCas4-dsh-V5-6myc

The V5-6myc fragment, flanked by Xba1 sites was generated by touch down PCR using pUAST-Rab35-6myc (Addgene #53503) as template.

Forward primer: TCTAGAAGCGGCACAGGCTCTGGCGGCAAACCGATTCCGAACCCGCTGCTGGGCCTGGATAGCACCAGTGGTGGATCCACCATGGAGCAAAAGCTC

Reverse primer: TCTAGAATACCGGTGATTACAAGTCCTCTTC

pCas4-dsh from ref.^36^ was digested with Xba1 and the V5-6xmyc fragment appended. The resulting plasmid was transformed into *w^11^*^18^ flies to generate random genomic insertions and an X chromosome insertion was recombined onto a *dsh^v^*^26^ *f^36a^* chromosome.

#### FmiΔcad

Generation of FmiΔcad, under control of the armadillo promoter, was described in (^69^). Structural analysis of the mammalian Fmi homolog CELSR2 showed that the extracellular domains form an antihelical homodimer involving the first 8 of 9 cadherin domains (the ones deleted in FmiΔcad) and that interaction is significantly diminished in constructs shorter than 5 repeats^31^. Therefore, removing domains 1-8, as in FmiΔcad, is expected to entirely eliminate the possibility of homodimerization in vivo and therefore communication and adhesion between cells. Functionally, we observe no ability to rescue the null phenotype, consistent with a complete lack in intercellular communication. FmiΔcad does, however, rescue cell competition^69^, and it elicits phenotypes that confirm it interacts intracellularly with other PCP components (this manuscript).

### Pupal wing dissections and immunohistochemistry

White prepupae were selected, raised at 25°C and aged to 28hr APF unless otherwise noted. For experiments involving Dsh mutants, only hemizygous male pupae were selected to ensure that the mutant *dsh* allele was the only allele present. All other experiments involved a combination of roughly equal male and female animals and the data were not disaggregated. Pupae were dissected from their pupal casing, affixed to double sided tape on a slide and were directly whole animal fixed for 30 min in 4% paraformaldehyde at room temperature. Pupal wings were then dissected from fixed pupae and moved to PBST (PBS with 0.1% Triton X-100). Primary antibody incubations (1:200 dilution) were performed overnight at 4°C and then washed in PBST for a total of 15 minutes. Incubations with secondary antibodies (and phalloidin if noted) were then performed for an hour at room temperature in PBST. Wings were then washed in PBST for a total of 15 minutes. Stained wings were mounted using 12 μl Vectashield mounting medium (Vector Laboratories). Primary antibodies were as follows: mouse monoclonal anti-V5 (1:200 dilution, Thermo-fisher, R960-25), guinea pig polyclonal anti-Pk[C] (1:800 dilution, (^60^), mouse monoclonal anti-Fmi (1:200 dilution, DSHB), rat monoclonal anti::HA (clone 3F10, 1:200 dilution, Roche), rat monoclonal anti-dEcad (1:200 dilution, DSHB), rabbit polyclonal anti-dsh (1:200)^43^, mouse monoclonal anti-Fz (1:200 DSHB 1C11), and mouse monoclonal anti-Myc (Thermo-fischer 1:200 sc-40). Secondary antibodies from Thermo Fisher Scientific were as follows: 488-donkey anti-mouse, 488-goat anti-rabbit, 596-donkey anti-mouse, 633-goat anti-guinea pig, Alexa 350 conjugated phalloidin was obtained from Thermo Fisher Scientific. Validations of these antibodies can be found in the product sheets from each vendor or in the studies in which they were generated.

Note that the anti-dsh antibody^43^ has insufficient sensitivity to detect endogenous levels of Dsh. In the experiments in Figures 2 and 4, one would expect to detect little signal outside of the N-Cad::Fz clones in any case as the surrounding tissue is mutant for *fz*. The anti-fz antibody (1:200 DSHB 1C11) is also insufficiently sensitive to detect endogenous levels of Fz. In the experiment in Figure 2F, no signal is detected outside the clone because only endogenous Fz, at a level below detection threshold, is present.

### Confocal microscopy and quantification

Immunostained pupal wing images were taken with a Leica TCS SP8 AOBS confocal microscope and processed in Fiji (version: 2.16.0/1.54g build:26d66057dd; Imagej). Adult wings were dissected and washed with 70% EtOH and mounted in DPX (Sigma) solution. All adult wings were imaged on a Nikon Eclipse E1000M equipped with a Spot Flex camera (Model15.2 64 MP).

Quantification of relative mean fluorescence intensity of staining at specified cell junctions was performed by tracing the junctions with a line of width 1 pixel drawn through the middle of the membrane, using the tracing and measurement tools built into the image processing software Fiji, reporting the average intensity along 8-10 cell junctions (approximately 40μm) of either the clone boundary, the interior of the clone, or at a distance away from the clone. Each measurement, corresponding to a single dot in the graphs, is taken from one clone per wing. Quantification of cytoplasmic levels (Fig. 3) was performed similarly, except intensity within an area in the cell interior rather than the cell membrane was measured.

The polarization score (Fig. 1J) was defined as the ratio of averaged fluorescence intensity on the bright sides over the averaged intensity on the opposite sides, where only sides on clone boundaries were scored. Each dot represents the value for an individual clone. A score equal to 1 is not polarized, and greater than 1 is polarized.

TIRF imaging was performed as described in ref. ^34^.

All wing images are shown with proximal to the left and anterior to the top.

All plasmids and fly strains will be available from the corresponding author or from the Bloomington Drosophila Stock Center (NIH P40OD018537).

## Data Availability Statement

All data are included in the manuscript, figures, and source data are present in the Source Data File.

## Acknowledgements

We would like to thank members of the Axelrod lab for fruitful discussions. We thank James Ferrell for fruitful discussions on feedback. Stocks obtained from the Bloomington Drosophila Stock Center (NIH P40OD018537) were used in this study. This work was funded by NIH R35GM131914 and R01HL16929901 (JDA). SBN was supported by the Novo Nordisk Foundation (grant awards NNF20OC0059462 and NNF21CC0073729) and the Stanford Bio-X Program. We acknowledge Elizabeth Weiner for suggesting the term ‘Offline,’ without whose linguistic ingenuity this work might lack its defining terminology and without whose due credit, the first author might face a more formidable review process at home than in the peer review process. The funders had no role in study design, data collection and analysis, decision to publish, or preparation of the manuscript.

## Author Contributions

JDA and ATW conceived the project. ATW helped build reagents and tools, performed experiments and wrote the original draft of the manuscript. KS designed and generated tools. SBN and BC helped design the project and performed experiments. GP-L performed experiments. JDA is the lead contact, supervised the project and wrote iterations of the manuscript. All authors contributed to and approved the final version of the manuscript.

## Supplementary Information

### Inventory of Supporting Information

**Supplementary Fig. 1**

**Supplementary Fig. 2**

**Supplementary Fig. 3**

**Supplementary Fig. 4**

**Supplementary Fig. 5**

**Supplementary Fig. 6**

**Source Data File** included as a separate Excel document

**Supplementary Figure 1.**
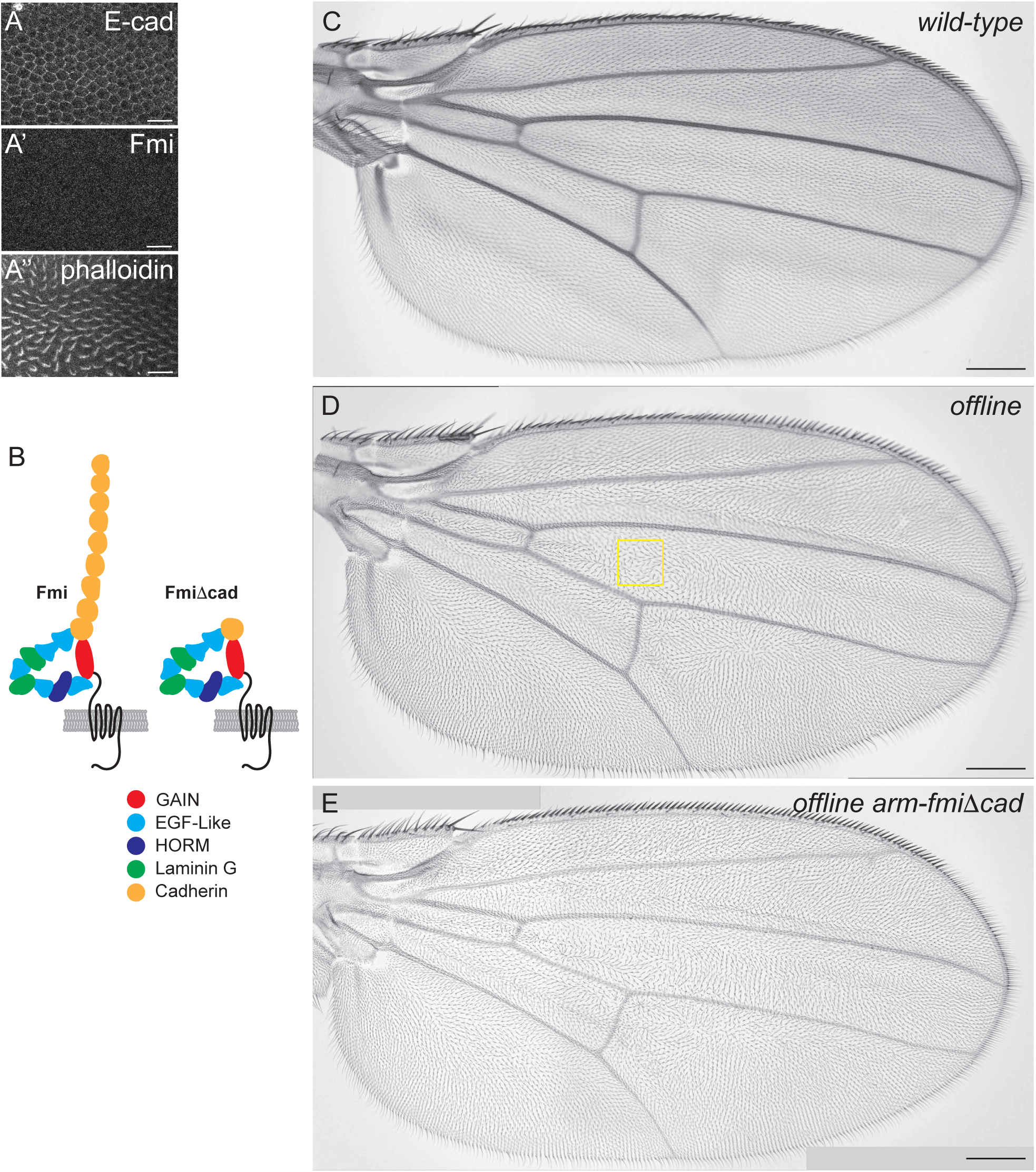
Validation of the offline system. A-A’’. Region of a *fmi^null^* offline (yellow box in B, QF2-1407, *fmi^E^*^59^ / *fmi^E^*^45^; QUAS-fmi (F17)) wing stained for E-Cadherin (A), Fmi (A’) and phalloidin (A’’), demonstrating no detectable Fmi. Scale bar = 10 μm. B. Cartoons of full length Fmi and FmiΔcad with conserved extracellular domains labeled, showing deletion of cadherin domains 1-8 in FmiΔcad. C. Adult wild-type wing showing uniform distally oriented hairs. Scale bar = 100 μm. D. Adult offline (*fmi^null^*) wing showing the stereotypical swirls and discontinuities characteristic of the core PCP mutant phenotype. The box near the anterior crossvein in the region of the swirl corresponds to the region of the image in Figure 1G. Scale bar = 100 μm. E. Adult offline (*fmi^null^*) arm-fmiΔcad wing showing the same stereotypical swirls and discontinuities characteristic of the core PCP mutant phenotype. Scale bar = 100 μm.

**Supplementary Figure 2.**
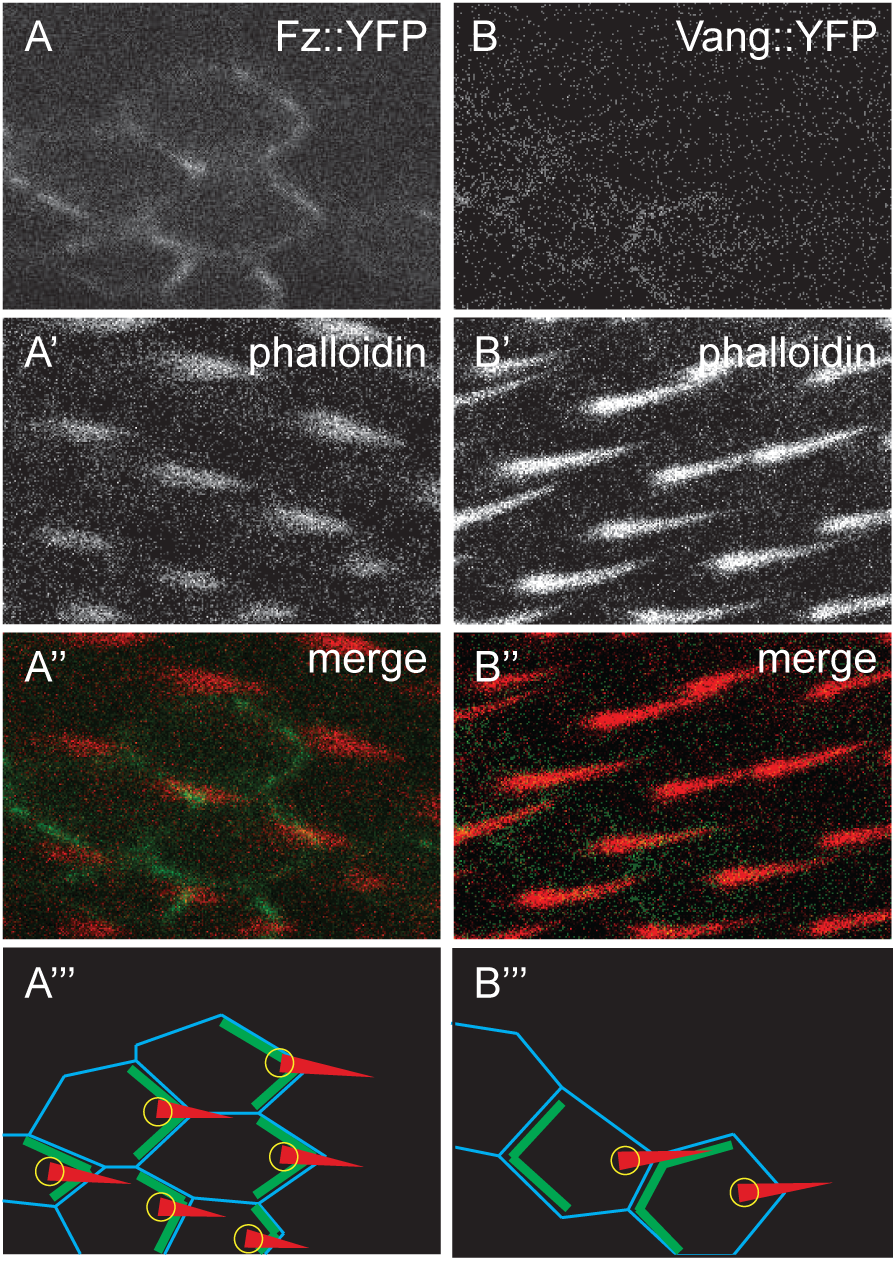
Prehair polarity in Fz::YFP and Vang::YFP Clones. A-A’’’. Single channel and merged images from 34hr APF pupal wings carrying Fz::YFP clones in a wild-type background, stained with phalloidin to reveal prehairs. Drawings are provided to clarify where prehairs emerge from the cells. B-B’’’. Single channel and merged images from 34hr APF pupal wings carrying Vang::YFP clones in a wild-type background, stained with phalloidin to reveal prehairs. Drawings are provided to clarify where prehairs emerge from the cells

**Supplementary Figure 3.**
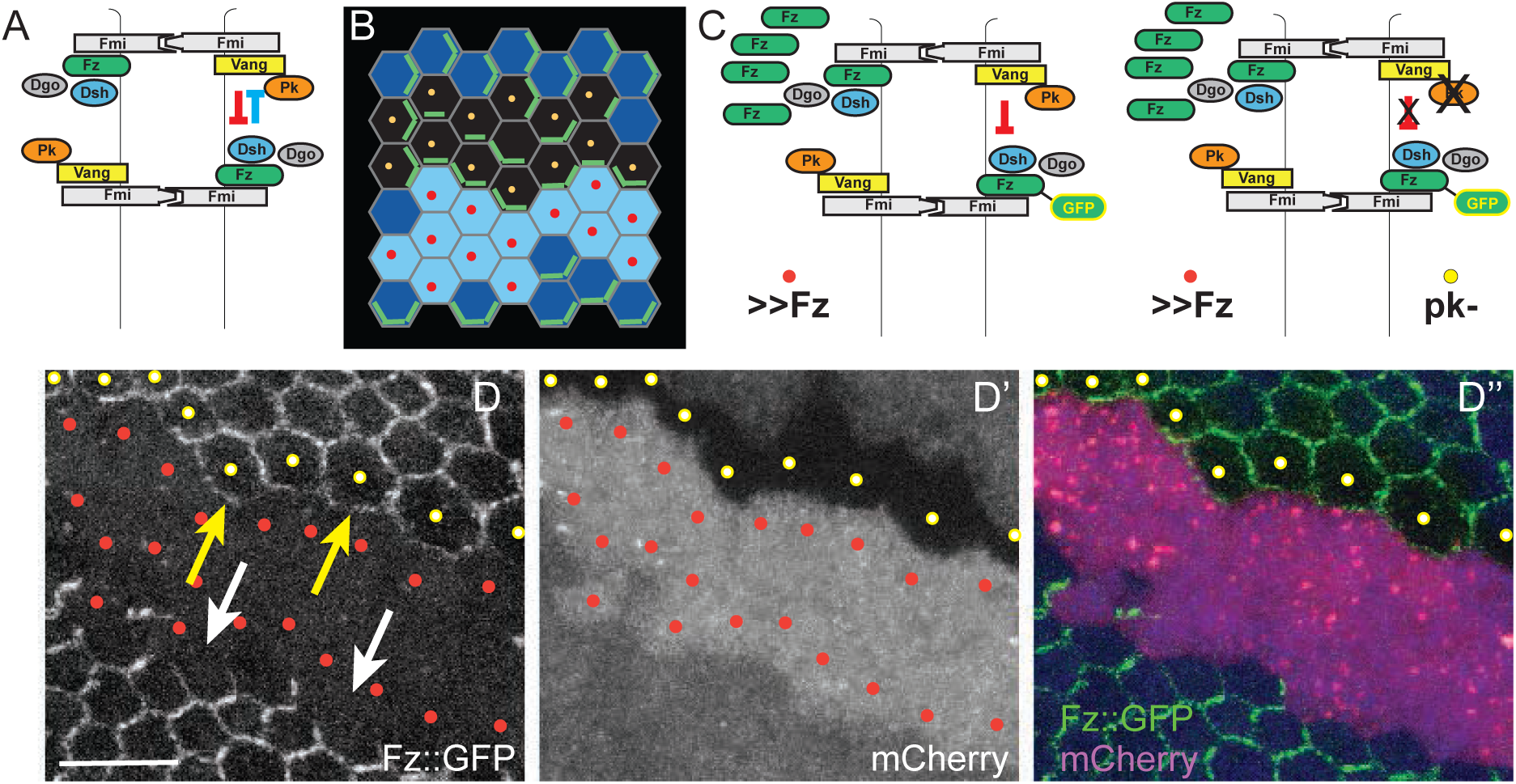
Fz overexpression excludes Fz from the boundary of neighboring cells. A. Cartoon showing Vang-dependent exclusion of Fz-Dsh complexes (red arrow) and Fz-dependent exclusion of Vang-Pk complexes (blue arrow). B. Schematic of reverse MARCM clones; red dots indicate the border of the Fz overexpressing clone (in D-D’’, red mCherry expressing cells) and yellow dots indicate *pk* mutant clonal cells facing Fz overexpressing cells (as in D-D’’). C. Schematics of MARCM experiment, showing Fz overexpressing cells recruiting Vang and excluding Fz from the boundary of the neighboring cell, and that this does not occur in neighbors that lack Pk. D-D’’. Fz overexpression excludes Fz::GFP from the membrane in neighboring wildtype cells (white arrows), but fails to exclude Fz::GFP in *pk* mutant cells (yellow dots; yellow arrows indicate membranous Fz::GFP facing Fz overexpressing cells in *pk* mutant cells). 26hr APF wing tissues. Scale bar: 10μm.

**Supplementary Figure 4.**
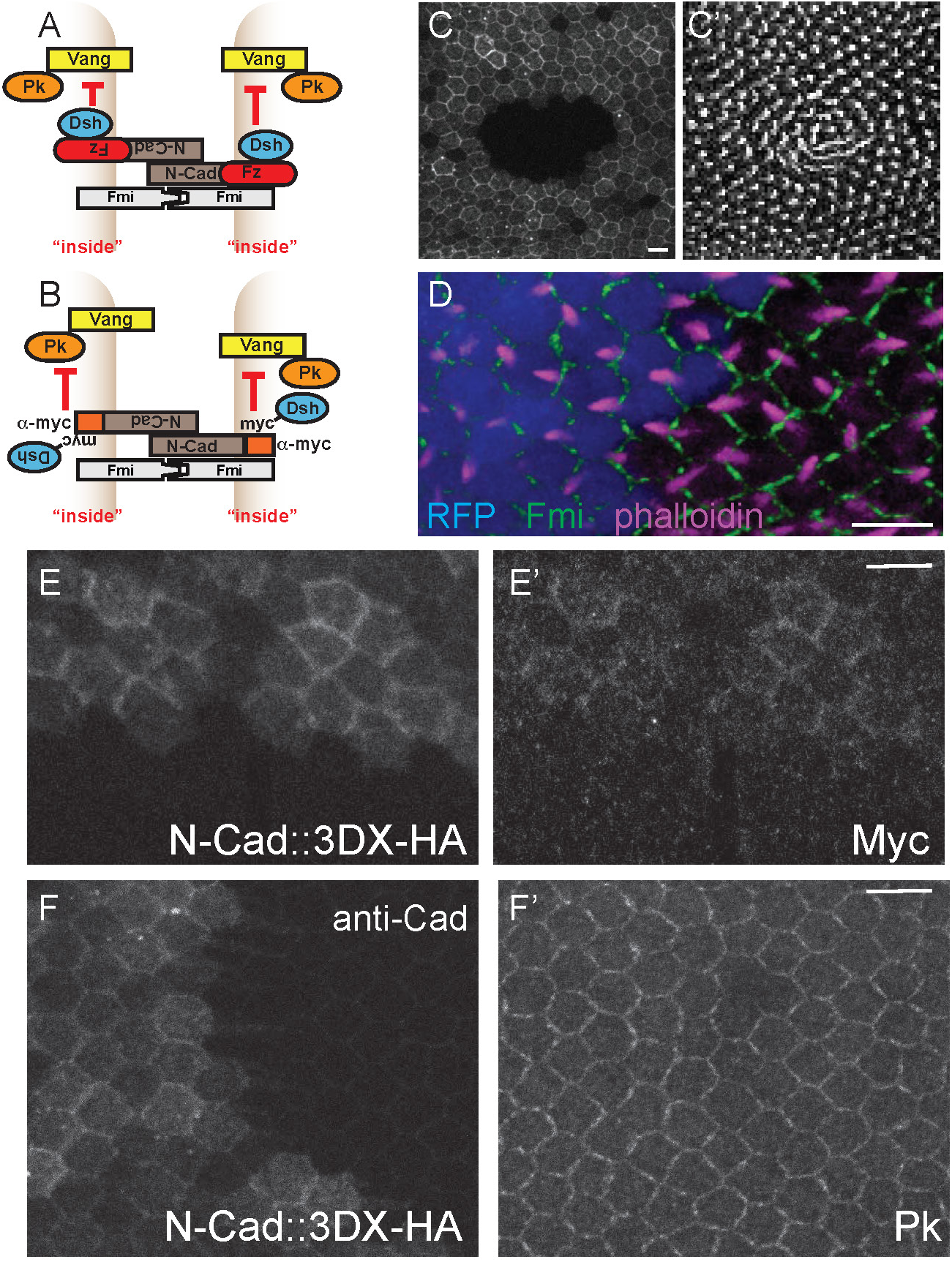
Validation of the Velcro system. A. Schematic of the Fz Velcro exclusion assay. B. Schematic of the Dsh Velcro exclusion assay. C,C’. A large clone expressing Fz Velcro wrapping around a region lacking Velcro, stained for V5 (N-Cad::V5-Fz; C) and phalloidin (C’). Polarity is perturbed by the clone boundary. D. Portion of a Dsh Velcro clone marked with RFP (blue), stained for Fmi (green) and phalloidin (magenta), showing prehairs of reversed polarity near the clone boundary. This is an example of a strong phenotype for Dsh Velcro (hsFLP; *dsh^V^*^26^, dsh-myc; act>>GAL4, UAS-RFP / UAS-N-Cad::HA-3DX). Many clones altered hair polarity only weakly or not at all. E,E’. N-Cad::HA-3DX expressed without Dsh-myc. Staining for HA (E) and with anti-myc reveals that endogenous Myc is recruited. F,F’. N-Cad::HA-3DX expressed alone, stained for anti-N-Cadherin (F) and for Pk (F’), demonstrating that N-Cad::HA-3DX itself does not disturb Pk localization. Note that the anti-N-Cadherin antibody detects Cadherin repeats from both N-Cad and endogenous E-Cad. E-Cad is faintly visible outside the N-Cad expression region. Scale bars = 5 μm.

**Supplementary Figure 5.**
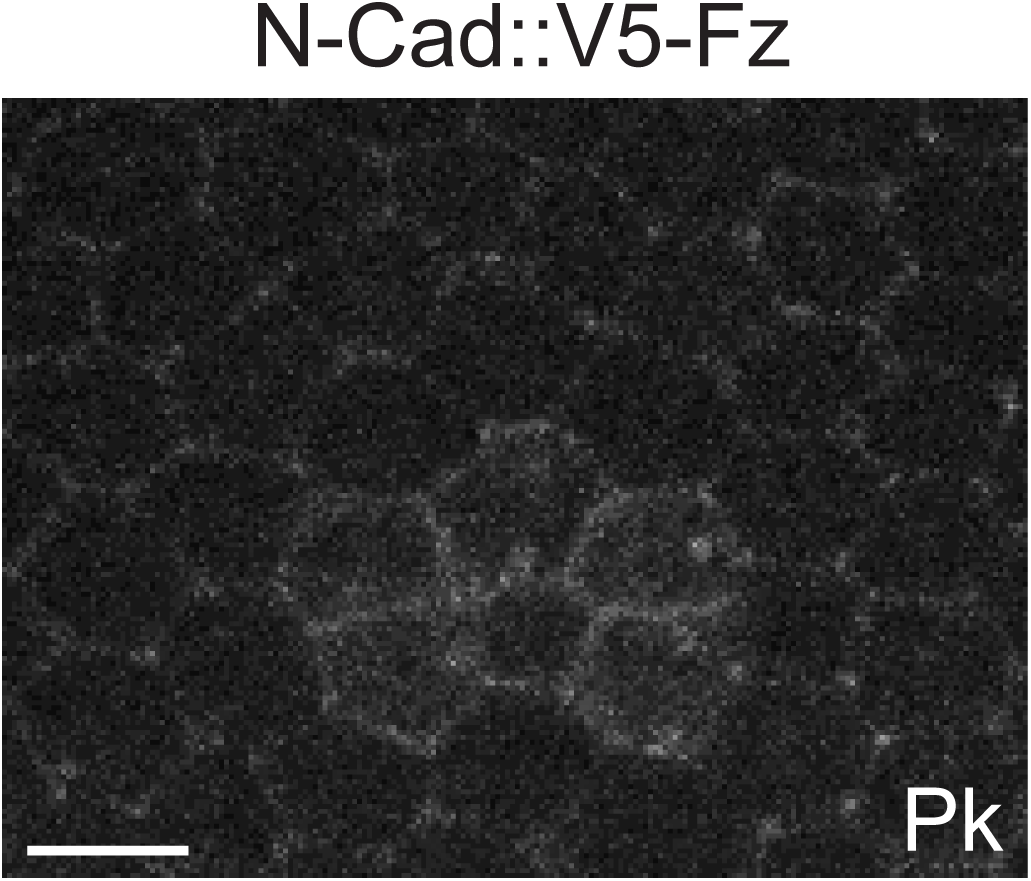
Control for Figure 4 showing Pk localizes to Fz Velcro clones in a *dsh^G63D^* mutant background. N-Cad::V5-Fz clones in *fz^null^*, *fmi^null^* background (Velcroed Fz, offline) and not stained for V5. Pk staining labels the clone indicating the signal indicating Pk recruitment is not a result of bleed through in Figure 4. Scale bar = 5 μm.

**Supplementary Figure 6.**
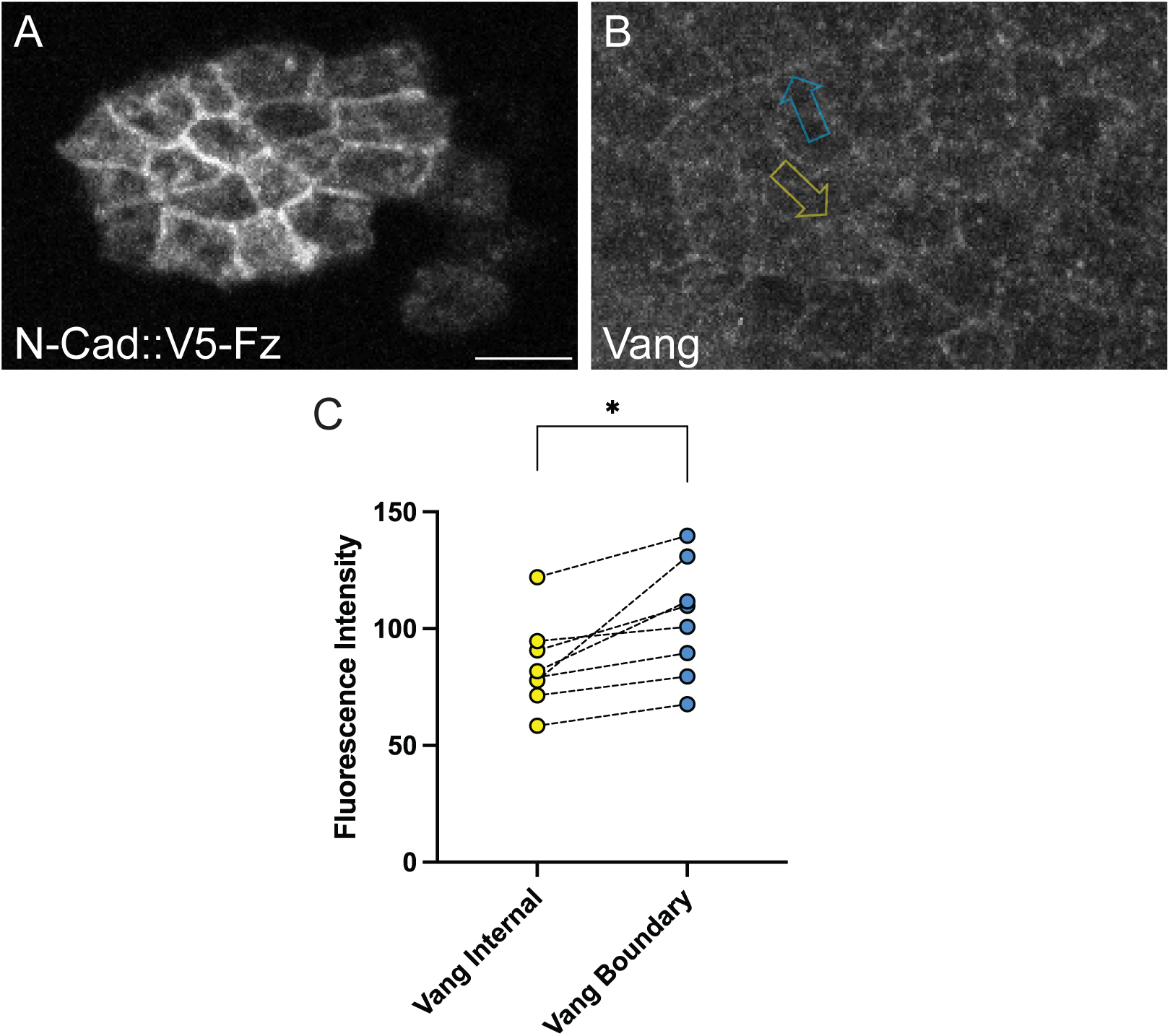
Vang Exclusion by Fz Velcro. A-B. N-Cad::V5-Fz clones in *fz^null^*, *fmi^null^* background (Velcroed Fz, offline) stained for V5 and Vang. N=8 individual wings from different animals C. Quantification of Vang fluorescence mean intensity linescans for paired data along internal N-Cad::V5-Fz contacts (yellow arrow in B) and clonal boundaries (blue arrow in B). Lines connecting data points denote the comparison from the an individual clone, each from a separate wing. Significance was tested with a two-tailed students t-test to compare means with p = .037 (*). Intensities at internal cell boundaries are lower than those at the clone periphery, indicating partial exclusion of Vang. Scale bar = 5 μm.

